# Mapler: Assessing assembly quality in taxonomically rich metagenomes sequenced with HiFi reads

**DOI:** 10.1101/2025.03.10.641994

**Authors:** Nicolas Maurice, Claire Lemaitre, Riccardo Vicedomini, Clémence Frioux

## Abstract

**Summary:** Metagenome assembly seeks to reconstruct the most high-quality genomes from sequencing data of microbial ecosystems. Despite technological advancements that facilitate assembly, such as Hi-Fi long reads, the process remains challenging in complex environmental samples consisting of hundreds to thousands of populations. Mapler is a metagenome assembly and evaluation pipeline with a focus on evaluating the quality of Hi-Fi long read metagenome assemblies. It incorporates several state-of-the-art metrics, as well as novel metrics assessing the diversity that remains uncaptured by the assembly process. Mapler facilitates the comparison of assembly strategies and helps identify methodological bottlenecks that hinder genome reconstruction.

**Availability and Implementation:** Mapler is open source and publicly available under the AGPL-3.0 licence at https://github.com/Nimauric/Mapler. Source code is implemented in Python and Bash as a Snakemake pipeline.

**Supplementary information:** Available online.

## 1. Introduction

Evaluating the quality of metagenome assemblies can be a challenging task, especially when no reference genome is available and when comparing samples with varying taxonomic richness and sequencing depths. Taxonomic richness refers to the number of distinct populations within the sample: microbial communities may consist of only a handful of populations, as in acid mine drainage communities [19], or of up thousands of distinct populations, as observed in soil ecosystems [18]. Assembly of metagenomic reads leads, in the best case scenario, to the reconstruction of genomes, but in most cases, to the generation of sequences of varying length called *contigs*. Those are then grouped into *bins*, presumed to originate from the same microbial populations; bins of sufficient quality are referred as *Metagenome-Assembled Genomes* (MAGs) [4]. A high-quality metagenome assembly is not only expected to yield high-quality bins, but also to be representative of the majority of the read sequences. Recent studies showed significant improvements in both the number and quality of bins obtained using highly accurate PacBio HiFi long reads [1]. However, in highly taxonomically rich ecosystems, assembly methods still struggle to reconstruct the numerous low-abundance genomes [1, 21], and it remains unclear how much of the sample these bins are representative of, resulting in a need for comparison and development of dedicated evaluation methods.

Several tools and pipelines exist to evaluate metagenomes. CheckM2 [6] assesses binned contigs based on the presence of marker genes, allowing the identification of MAGs from bins with low contamination and high completeness scores. The PacBio HiFi-MAG-Pipeline [16] is a pipeline developed to identify high-quality MAGs from previously generated metagenome assemblies. It follows a “completeness-aware” strategy based on CheckM2 and several state-of-the-art binning tools, incorporating stringent filtering criteria to exclude low-quality bins, which are common in taxonomically rich ecosystems. MetaQUAST [13] performs a reference-based evaluation, either using user-defined references or retrieving references via taxonomic assignment. However, in complex ecosystems, many species are absent from databases, thus limiting its effectiveness. Finally, custom metrics or visualizations have also been employed for method validation [1, 7]. For example, the percentage of reads aligned to the assembly has been used to validate metagenome assembly in [1]. Nevertheless, these approaches are rarely documented nor provided in an easy-to-use implementation that allows for replication on new datasets.

In this work, we present Mapler, a metagenomic assembly and evaluation pipeline. It avoids filtering out any sequences, it does not rely on the availability of reference sequences, and it considers both unassembled reads and unbinned contigs. Mapler integrates several state-of-the-art tools as well as novel metrics and visualizations based on read-to-contig alignments. It provides a broad view of the sequence characteristics after assembly and binning, in order to identify the bottlenecks faced during bioinformatic processes. Mapler is therefore an effective way to examine assembly in taxonomically rich ecosystems, where high-quality bins and references are scarce.

## 2. Software description

### 2.1. Pipeline

Mapler is a Snakemake [14] pipeline dedicated to the evaluation of taxonomically rich metagenome assemblies of HiFi long reads. It can be run either locally or on Slurm-based computing environments. Its modular design allows for easy integration of additional custom steps in the pipeline or modification of existing ones. The pipeline can run multiple steps in parallel, including analysing multiple samples at once. The structure of the pipeline is illustrated in Figure 1A.

**Figure 1:**
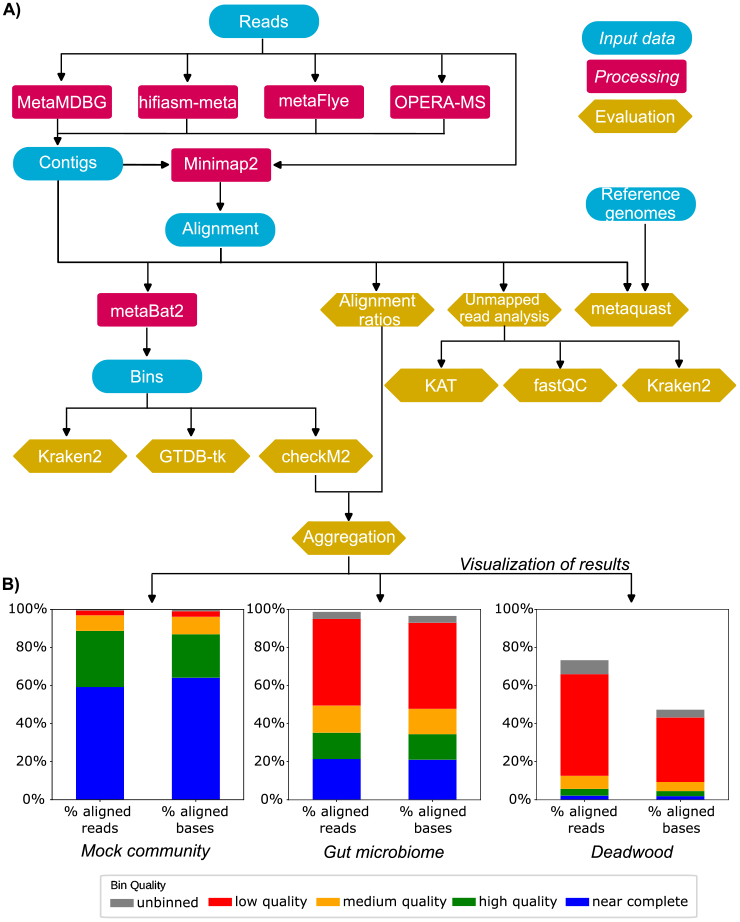
(A) Overview of the Mapler pipeline. Contigs and bins can either be generated by the pipeline or given as input. Several long-read assemblers are integrated in Mapler. (B) Example of Mapler’s output. Histograms show the aligned read/base percentages for bins of different quality and reveal the increasing complexity of different ecosystems, from the mock community to the gut microbiome sample and the highly diverse deadwood sample, all three assembled with metaMDBG.

### 2.2. Integrated tools

While its focus being on evaluation, Mapler integrates state-of-the-art tools for assembly and binning suitable for HiFi sequencing data: metaMDBG [1], hifiasm-meta [8], metaFlye [10], OPERA-MS [2], and MetaBAT 2 [9]. Users may alternatively skip the assembly and/or binning steps by providing their own input contigs and/or bins. Each bin can be taxonomically classified using either GTDB-Tk [5], or Kraken 2 [20] in order to facilitate the comparison with the taxonomic assignment of the reads. By default, bins are qualitatively assessed with CheckM2 [6] and categorized according to the following levels of completeness (comp.) and contamination (cont.): near complete (single contig, ≥99% comp., ≤1% cont.), high quality (≥90% comp., ≤5% cont.), medium quality (≥50% comp., ≤10% cont.), and low quality for the remaining bins. These criteria match the completeness and contamination estimates used by the Genomic Standards Consortium [3] for defining low-quality to high-quality MAGs. MetaQUAST [13] is also integrated to compare contigs with reference genomes, if available and provided as input by the user.

### 2.3. Novel metrics

Mapler aligns the reads on the contigs with Minimap2 [11], and uses these alignments to calculate various metrics. The *aligned read percentage* is the number of reads aligned to at least one contig divided by the total number of reads, while the *aligned base percentage* is the number of read bases aligned to at least one contig, divided by the total length of the reads. These metrics can be computed with or without the binning information. In the former case, the percentage is separately calculated for reads or bases that align to contigs belonging to bins of near complete, high, medium or low quality, or to contigs that were assembled but not binned. A text report is produced for both the binning-aware and binning-unaware versions, and a summarizing plot is generated for the binning-aware version (Figure 1.B). In cases where a read is aligned to multiple contigs, it is only taken into account for the highest bin quality level.

Another analysis proposed by Mapler is the comparison of the sets of reads aligned or unaligned to the contigs, in order to gain insight into the characteristics of reads that participate in, or have been excluded from the assembly. Both sets are analyzed separately with the following tools:

- FastQC (https://github.com/s-andrews/FastQC), used to assess read quality and generate a comprehensive report. It can be used to check whether the assembly process is more effective on higher quality reads, longer reads, or reads with a certain GC ratio.
- K-mer Analysis Toolkit (KAT) [12], which computes the abundance of assembled and unassembled reads. The abundance of a given read is estimated by its median k-mer abundance, with k-mer abundances being computed from the full read dataset. Mapler integrates these results to visualize both distributions with two overlapping histograms.
- Kraken 2 [20], alongside Krona [15], is used to analyze the taxonomic composition and abundance of both sets of reads, providing insight on over- or under-represented clades in the assembly.

## 3. Application

We demonstrated Mapler’s ability to evaluate assemblies of diverse samples on three datasets of increasing taxonomic complexity, sequenced with PacBio Sequel II SMRT.

- *Mock community* : the ZymoBIOMICS Gut Microbiome Standard D6331 (SRR13128014) consists of 21 populations, including 17 species and 5 strains of *Escherichia coli*. The sample contains 18.0 Gbp spread over 1, 978, 852 reads.
- *Gut microbiome*: A pooled extraction of four stool samples from adult humans following a vegan diet (SRR15275211). Human digestive microbiomes generally host a few hundreds of species. The sample contains 18.8 Gbp spread over 1, 904, 159 reads.
- *Deadwood* : four separately sequenced samples of deadwood that were co-assembled as in [17]. The samples (SRR28211698 to SRR28211701) contain a total of 16.1 Gbp spread over 866, 007 reads.

Each dataset was processed by Mapler with metaMDBG, hifiasm-meta, and metaFlye.

Mapler first summarises in scatter plots the bins obtained in each sample (Supp. Fig. 1), highlighting that the number and quality of bins vary across the datasets. Compared to the *Mock community*, the number of low quality bins is much higher in the *Gut community* and *Deadwood* samples, due to either low completeness or high contamination scores.

Mapler then generates, after mapping reads to contigs and bins, plots that highlight a decreasing proportion of reads assembled and binned at each quality level as dataset complexity increases (Fig. 1B). More precisely, on the metaMDBG assemblies, 96.9% of reads and 96.1% of bases map to bins of at least medium quality in the *Mock community*, while in the *Gut microbiome* these values drop to 49.5% and 47.8%, respectively. Furthermore, *Deadwood*’s high diversity and lower sequencing depth result in a lower-quality assembly with only 12.6% of reads and 9.4% of bases aligned with bins of medium quality or higher.

Because a significant proportion of reads of the *Deadwood* sample did not participate in the assembly (26.7% of reads and 52.7% of bases did not align with any contig), we compared the aligned and unaligned reads in this sample. Despite the read length variation in the original sample, assembled and unassembled reads are of similar length (18, 437 and 18, 583 base pairs on average, respectively, see Supp. Fig. 2). Taxonomic assignment of reads with Mapler illustrates that some microbial populations were only detected in unassembled reads, such as several species of *Legionella* (Supp. Fig. 3). Sequences were also generally assigned with less precision in the unassembled reads (56% of bacteria are assigned at the phylum level in the unassembled reads, compared to 76% in the assembled reads), suggesting that most low-abundant populations remain unknown in databases. As expected, unassembled reads were mostly made up of rare k-mers: nearly all unassembled reads have a median k-mer abundance as low as 1 (Supp. Fig. 4). These results suggest that assemblers and binners used in the metagenome analysis could not improve the results by much, and that a deeper sequencing would rather be needed to enhance the quality of the assembly. We nonetheless compared the *Deadwood* assemblies performed with different metagenome assembly tools. MetaMDBG outperformed the other assemblers in terms of total captured diversity: 52.7% of bases do not align with any contig, compared to 76.7% for metaFlye and 66.1% for hifiasm-meta (Supp. Fig. 5). Conversely, hifiasm-meta outperformed metaMDBG in term of bases aligned to at least medium-quality bins (14.2% versus 12.6%).

We recorded the execution time of the pipeline on the three datasets. For each dataset, we executed the pipeline on a Intel(R) Xeon(R) CPU E5-2670 v3 @ 2.30GHz node, allocating a total of 48 CPUs and 200G of memory. The detailed breakdown of how much memory was allocated to each substep of the pipeline is described in Supplementary Table 1. We performed the evaluations on the three samples separately. For each sample, we evaluated the time it took to evaluate the assembly and binning quality of three assemblers (the assembly and binning was performed separately). The analysis of the *Mock community* took 2 hours and 4 minutes in wallclock time to run, followed by the *Gut microbiome* with 3 hours and 46 minutes and, finally, the *Deadwood* with 5 hours and 3 minutes.

## 4. Conclusion

Mapler is a metagenome evaluation pipeline that allows a thorough examination of assembly and binning, implemented in an easy-to-use and customizable workflow. Mapler is specifically implemented to analyze HiFi long-read datasets that are currently the most suitable to characterize taxonomically rich microbial ecosystems. On top of integrating multiple state-of-the-art evaluation methods, Mapler incorporates new evaluation metrics such as the aligned read and aligned base percentages. When combined with the bin quality information, these metrics provide a way to measure how much of the sample’s original diversity was assembled at each level of quality, and highlight potential assembly issues. In cases where a significant proportion of reads cannot be aligned back to the assembly, comparing the assembled and unassembled reads provides further insight into the reasons why the assembly may not be sufficiently representative of the sample, and whether the contigs are missing key taxa that are only present in the reads.

## 6. Author contributions statement

CF, CL, NM and RV conceived the experiments; NM conducted the experiments; NM developed the software; NM, CF, CL and RV tested the software; CF, CL, NM and RV wrote the manuscript; All authors read and approved the manuscript.

## 7. Acknowledgments

This work was supported by the French National Research Agency (ANR) France 2030 PEPR Agroécologie et Numérique MISTIC ANR-22-PEAE-0011. We acknowledge the GenOuest bioinformatics core facility (https://www.genouest.org) for providing the computing infrastructure. A CC-BY public copyright license (https://creativecommons.org/licenses/by/4.0/) has been applied by the authors to the present document, in accordance with the grant’s open access conditions.

