## Supplementary Material for "Mapler: Assessing assembly quality in taxonomically rich metagenomes sequenced with HiFi reads"

Nicolas Maurice[1,2], Claire Lemaitre[1], Riccardo Vicedomini[1] and Clémence Frioux[2]
[1]GenScale, Univ. Rennes, Inria, CNRS UMR 6074, Rennes, F-35000, France
[2]Inria, Univ. Bordeaux, INRAE, F-33400 Talence, France

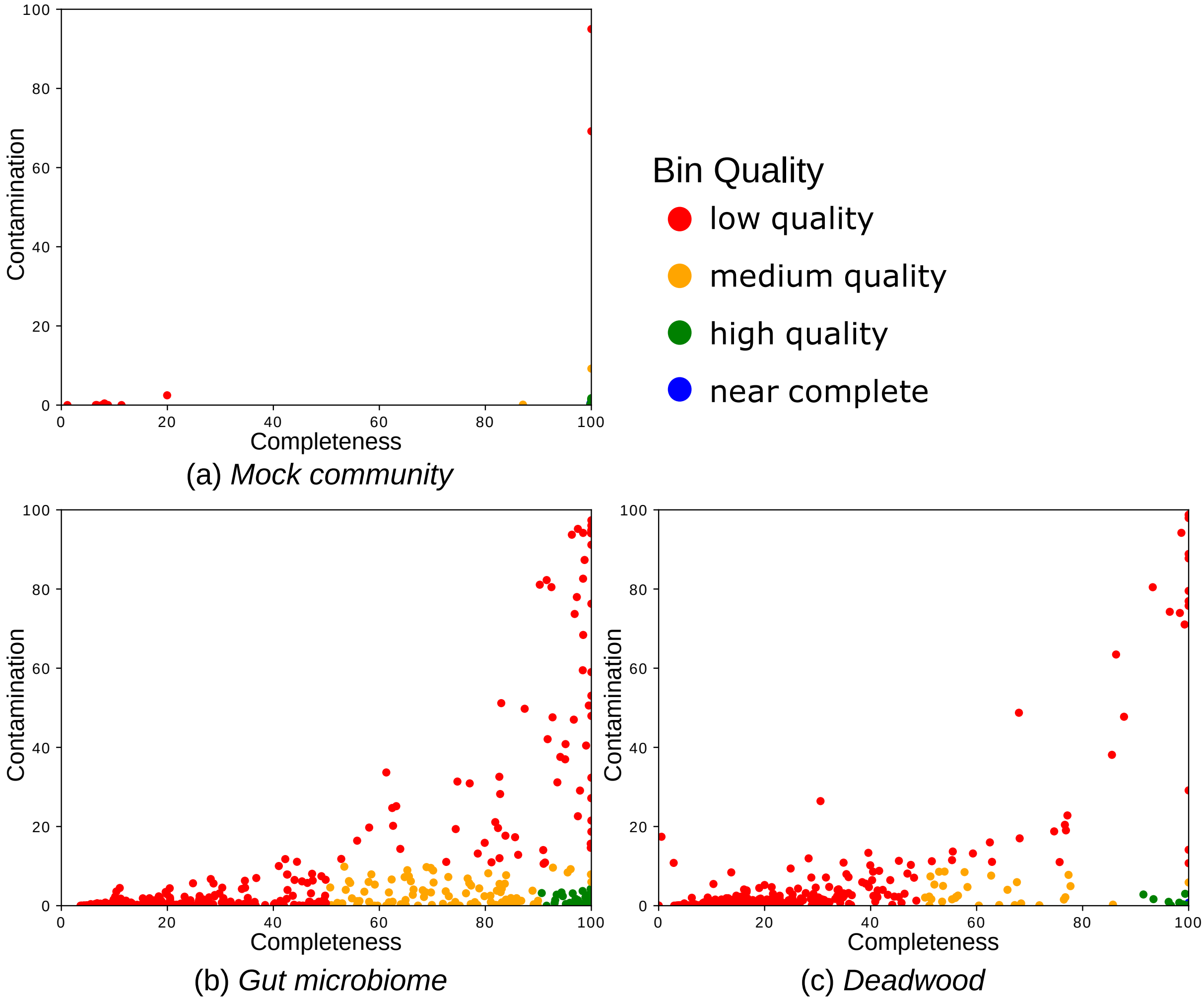


Figure S1: Comparison of the binning results between three samples, (a) mock community, (b) Gut microbiome, and (c) Deadwood, each assembled with metaMDBG. Green, orange and red dots denote the quality of the bin: high, medium and low respectively.

| Rule | Threads | Memory (MB) | Time limit (hours) |
|---|---|---|---|
| metaquast | 1 | 20,000 | 24 |
| read_contig_mapping_evaluation | 1 | 10,000 | 2 |
| checkm | 16 | 50,000 | 24 |
| checkm_report_writer | 1 | 50,000 | 1 |
| kraken2_on_bins | 12 | 100,000 | 4 |
| gtdbtk_on_bins | 20 | 50,000 | 8 |
| read_contig_mapping_plot | 1 | 30,000 | 2 |
| extract_unmapped_reads | 24 | 50,000 | 24 |
| kraken2 | 12 | 100,000 | 4 |
| kat_sect | 12 | 200,000 | 4 |
| fastqc | 1 | 5,000 | 2 |

Table S1: Resource allocation for each snakemake evaluation rule used for the time evaluation of all analyses.

FastQC Report

### Basic Statistics

| Measure | Value |
|---|---|
| Filename | mapped_reads.fastq |
| File type | Conventional base calls |
| Encoding | Sanger / Illumina 1.9 |
| Total Sequences | 634609 |
| Total Bases | 11.7 Gbp |
| Sequences flagged as poor quality | 0 |
| Sequence length | 1006-50047 |
| %GC | 59 |

| Measure | Value |
|---|---|
| Filename | unmapped_reads.fastq |
| File type | Conventional base calls |
| Encoding | Sanger / Illumina 1.9 |
| Total Sequences | 231398 |
| Total Bases | 4.3 Gbp |
| Sequences flagged as poor quality | 0 |
| Sequence length | 1000-48288 |
| %GC | 56 |

Figure S2: Statistics of assembled reads (”mapped reads”) and non-assembled reads (”unmapped reads”)from the metaMDBG assembly of the *Deadwood* sample, reported by FastQC.

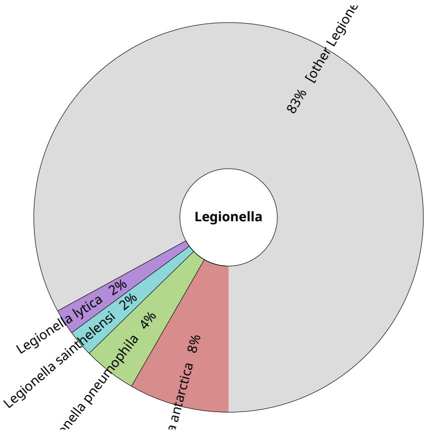


(a) Aligned *Legionella*-assigned reads

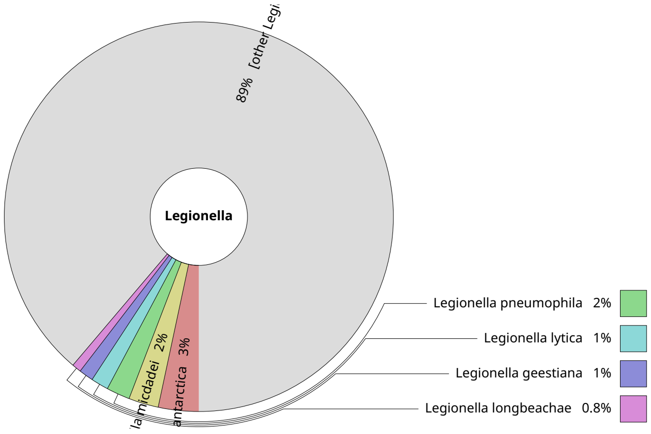


(b) Unaligned *Legionella*-assigned reads

Figure S3: Comparison of the taxonomic assessment of (a) aligned and (b) unaligned reads from the metaMDBG assembly of the *Deadwood* sample, with a focus on *Legionella* genus (0.05% of unaligned reads, 0.006% of aligned reads)

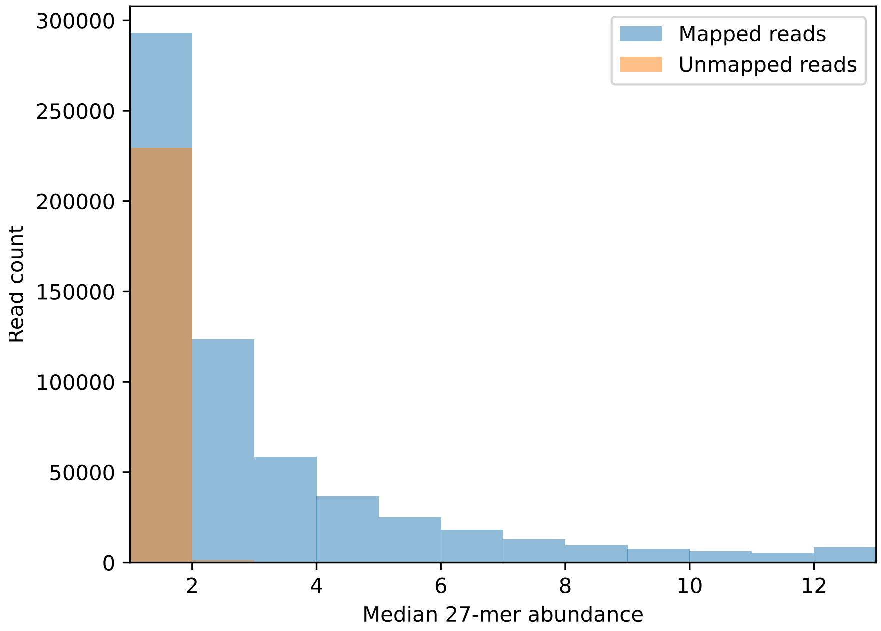


Figure S4: Comparative abundance of reads aligned (blue) or not aligned (orange, i.e. not assembled) to the metaMDBG assembly of the *Deadwood* sample

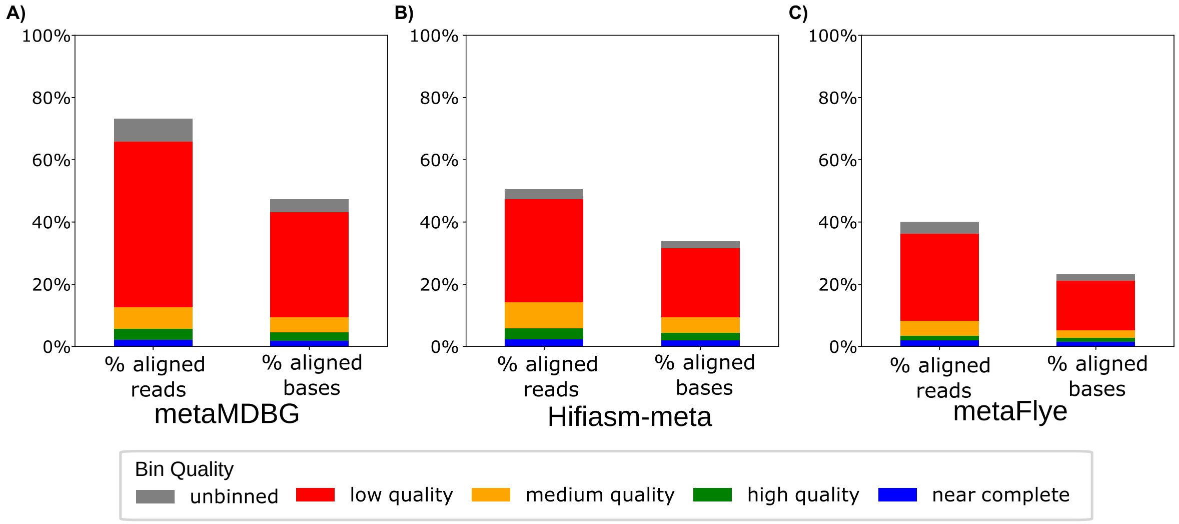


Figure S5: Comparison of the captured diversity for the *Deadwood* sample using three assemblers: (A) metaMDBG, (B) Hifiasm-meta, and (C) metaFlye.
